# Effects of *NT5C2* germline variants on 6-mecaptopurine metabolism in children with acute lymphoblastic leukemia

**DOI:** 10.1101/2020.10.13.338384

**Authors:** Chuang Jiang, Wenjian Yang, Takaya Moriyama, Chengcheng Liu, Colton Smith, Wentao Yang, Maoxiang Qian, Ziping Li, Morten Tulstrup, Kjeld Schmieglow, Kristine R. Crews, Hui Zhang, Ching-Hon Pui, William Evans, Mary Relling, Smita Bhatia, Jun J. Yang

## Abstract

6-mercaptopurine (6-MP) is widely used in the treatment of acute lymphoblastic leukemia (ALL), and its cytotoxicity is primarily mediated by thioguanine nucleotide metabolites (TGN). Recent genome-wide association study has identified germline polymorphisms (e.g., rs72846714) in the *NT5C2* gene associated with 6-MP metabolism in patients with ALL. However, the full spectrum of genetic variation in *NT5C2* is unclear and its impact on 6-MP drug activation has not been comprehensively examined. To this end, we performed targeted sequencing of *NT5C2* in 588 children with ALL and identified 121 single nucleotide polymorphisms (SNPs) nominally associated with erythrocyte TGN during 6-MP treatment (*P* < 0.05). Of these, 61 variants were validated in a replication cohort of 372 children with ALL. After considering linkage disequilibrium and multivariate analysis, we confirmed two clusters of variants, represented by rs72846714 and rs58700372, that independently affected 6-MP metabolism. Functional studies showed that rs58700372 directly altered the activity of an intronic enhancer, with the variant allele linked to higher transcription activity and reduced 6-MP metabolism (lower TGN). By contrast, rs72846714 was not located in a regulatory element and instead its association signal was explained by linkage disequilibrium with a proximal functional variant rs12256506 that activated *NT5C2* transcription in-*cis*. Our results indicated that *NT5C2* germline variation significantly contributes to inter-patient variability in thiopurine drug disposition.

## Introduction

6-mercaptopurine (6-MP) is widely used as an anticancer and immunosuppressive agent [1–8]. In acute lymphoblastic leukemia (ALL), prolonged daily 6-MP exposure during maintenance therapy is one of the major components of contemporary ALL treatment regimens [7, 9–13]. 6-MP has a narrow therapeutic index and frequent dose adjustment is needed to balance between antileukemic efficacy and treatment toxicity in patients [7, 13, 14]. However, large inter-patient variability in 6-MP disposition poses a significant challenge to clinically determine the optimal 6-MP dosage for each individual [15, 16].

As a prodrug, 6-MP is converted into thioguanine nucleotides TGN (thioguanosine monophosphate [TGMP], thioguanosine diphosphate [TGDP], thioguanosine triphosphate [TGTP]) via a series of enzymatic reactions [8, 17]. TGTP is further reduced to deoxythioguanosine triphosphate (TdGTP), which is then incorporated into double-stranded DNA (DNA-TG) to trigger futile mismatch repair and eventually apoptosis [7, 18–20]. Levels of these intracellular 6-MP metabolites can inform drug toxicity and response and have the potential to guide treatment individualization. For example, the NOPHO ALL2008 study demonstrated that DNA-TG concentration is significantly associated with the risk of leukemia relapse [21]. Variation in erythrocyte TGN level during maintenance therapy has been associated with relapse-free survival in some patients with ALL [12]. Therefore, therapeutic drug monitoring of thiopurine metabolites may be clinically relevant for patients receiving 6-MP.

6-MP metabolism is regulated by several key enzymes (e.g., NT5C2, TPMT, and NUDT15), and genetic variants in these genes have been linked to 6-MP toxicity and/or resistance. For example, inherited deficiency of TPMT and/or NUDT15 causes excessive accumulation of 6-MP active metabolites and severe toxicity, and genotype-guided dose reduction can dramatically mitigate side effects of this drug [22–27]. By contrast, somatic *NT5C2* mutations are common in relapsed ALL and encode hyperactive nucleotidase that degrades 6-MP active metabolites thioinosine monophosphate (TIMP) and TGMP, leading to lower TGTP, decreased DNA-TG, and 6-MP resistance [28–32]. The presence of sub-clonal ALL *NT5C2* mutations was associated with high risk of treatment failure and inferior outcome after relapse [33]. A recent genome-wide association study (GWAS) reported germline variants at the *NT5C2* locus (e.g., rs72846714) strongly associated with erythrocyte TGN level during ALL therapy [34].

Although the genome-wide significant association at the *NT5C2* locus signifies the impact of inherited variation in this gene on 6-MP pharmacokinetics, the full spectrum of genetic polymorphism in *NT5C2* is unclear and its impact on drug activation has not been systematically examined. To this end, we performed targeted sequencing of *NT5C2* to evaluate the contribution of germline *NT5C2* variants to inter-individual variations in 6-MP metabolites in children with ALL.

## Materials and methods

### Patients and therapy

The discovery cohort consisted of patients prospectively enrolled onto the Children’s Oncology Group (COG) AALL03N1 study. Details of enrollment criteria and study design have been described previously [16, 26]. Briefly, patients were treated on COG AALL03N1 frontline protocols for newly diagnosed ALL (or per frontline protocols for small number of patients), were enrolled onto this study for a total of six months maintenance therapy that included a daily dose of oral 6-MP. The protocol planned dose of 6-MP during maintenance phase was 75 mg/m^2^. per day for the COG AALL03N1 cohort, with provisions for dose adjustment based on the degree of myelosuppression (white blood cell count) and/or occurrence of infections.

The replication cohort consisted of children with ALL treated on the St Jude Children’s Research Hospital Total Therapy XV (St. Jude Total XV) protocol, with the planned dose of 75 mg/m^2^. in the low-risk arm or 50 mg/m^2^. in the standard- or high-risk arm during the phase of continuation therapy when TGN was measured [35]. *TPMT* and *NUDT15* genotyping was performed as described previously [26], and patients with known risk variants in either one of these two genes were excluded from the main analyses (N=74 and 47 in the COG AALL03N1 and St. Jude Total XV cohorts, respectively).

This study was approved by respective institutional review boards, and informed consent was obtained from parents, guardians, or patients, as appropriate.

### 6-MP metabolites measurement

Erythrocyte TGN levels were measured monthly for each participant during the six months study period of the COG AALL03N1 study, as described previously [16]. We generated a summarized TGN value for each patient based on their longitudinal TGN data using a linear mixed-effect model adjusting for time point of each measurement and prescribed 6-MP dose recorded monthly prior to the TGN measurement [26]. For St. Jude Total XV patients, TGN was measured at the start of Reinduction I during continuation therapy, using methods described previously [36, 37], and total 6-MP dose two weeks prior to sampling was recorded as total mg^2^/m^2^. TGN data from patients who received fewer than 12 of the 14 doses of 6-MP prior to drug metabolite measurement (N=18) or those with non-adherence based on self-report (N=4) were excluded.

### *NT5C2* targeted sequencing and SNP genotype imputation

Germline DNA was extracted from blood or bone marrow samples collected during clinical remission. Illumina dual-indexed libraries were created from genomic DNA and pooled in sets of 96 before hybridization with customized Roche NimbleGene SeqCap EZ probes (Roche, Roche NimbleGen, Madison, WI, USA) to capture the *NT5C2* genomic region. Quantitative PCR was used to define the appropriate capture product titer necessary to efficiently populate an Illumina HiSeq 2000 flow cell for paired-end 2 × 100 bp sequencing. Coverage of greater than 20 x depth was achieved across more than 80% of the targeted regions for nearly all samples. Sequence reads in the FASTQ format were mapped and aligned using the Burrows-Wheeler Aligner (BWA) and genetic variants were called using the GATK pipeline (version 3.1), as previously described [38–40], and annotated using the ANNOVA program with the annotation databases including RefSeq.

In addition to targeted sequencing, we also performed imputation for variants at the *NT5C2* locus, on the basis of SNP genotype data from Affymetrix SNP6 arrays of COG and St. Jude patients [26]. We employed a large reference panel of human haplotypes from the Haplotype Reference Consortium (HRC r1.1 2016) in Michigan Imputation Server with ShapeIT (v2.r790) as the phasing tool for imputation, following procedures described previously [41]. Imputed variants were filtered based on imputation quality, allele frequency, call rate, and deviation from Hardy-Weinberg equilibrium. Specifically, variants with one of the following features were excluded: imputation quality metric r^2^ less than 0.3 (indicating inadequate accuracy of the imputed genotype); minor allele frequency in less than 1% in either one of our two patient cohorts; Hardy-Weinberg equilibrium p value less than 5.00×10^-4^; variants located outside *NT5C2* locus (beyond 100kb of the first and last *NT5C2* exons).

### Statistical analyses

The association of *NT5C2* genotype with TGN was analyzed in patients with wildtype genotype in *TPMT* and *NUDT15* to avoid confounding effects. A linear regression model was used to test association, with 6-MP dose and genetic ancestry as covariates. R statistical software (version 3.5; http://www.r-project.org/) was used for all analyses unless indicated otherwise. Regional association plots were created by Locus zoom (http://locuszoom.org/). We also explored association of NT5C2 variants with TGN in patients with heterozygous genotype at known TPMT and/or NUDT15 variants.

### *NT5C2* recombinant protein production and 5’-nucleotidase activity assay

C-terminal Hisx6-tagged full-length cDNA for wildtype *NT5C2* was cloned into the pET30a expression vector using NdeI and HindIII restriction sites. Point mutations, p. Thr3Ala, p.Met15Thr and p.Ile528Val, (hereafter referred to as p.T3A, p.M15T, and p.I528V) were introduced by using the QuickChange II XL Site-Directed Mutagenesis Kit (Agilent Technologies). NT5C2 recombinant protein production and purification were performed using previously described methods [28]. Successful purification of NT5C2 proteins was confirmed by Coomassie brilliant blue staining (**Figure S1**). 5’-nucleotidase activity was measured by quantifying the release of phosphate from TIMP (Toronto Research Chemicals), or TGMP (Jena Bioscience), using the SensoLyte Malachite Green Assay kit (AnaSpec) in assay buffer (MgCl_2_ 2 mmol/L, NaCl 120 mmol/L, KCl 5 mmol/L, glucose 10 mmol/L, HEPES 20 mmol/L). NT5C2 protein (300 ng) was incubated with 40 mmol/L TIMP, or TGMP at 37°C for 30 minutes. Experiments were assessed in triplicates.

### Functional annotation and characterization of non-coding variants in *NT5C2*

Chromatin accessibility in human hematopoietic cells were extracted from GSE74912 [42]. Transcription factor ChIP-seq clusters and DNase hypersensitive site data were from ENCODE project [43, 44]. All peaks were aligned to GRCh37/hg19 on UCSC genome browser [45–47].

To functionally characterize potential function of non-coding variants in *NT5C2*, 356-600bp regions encompassing the SNP of interest were amplified using CloneAmp HiFi PCR Premix (Clontech), and then cloned into the pGL4.23-mini/P vector including a minimal SV40 promoter upstream of firefly luciferase. The variants were introduced by site-directed mutagenesis, by using the QuickChange Lightning Site-Directed Mutagenesis Kit (Agilent). Primers used in this experiment were listed in **Table S1**. Human erythroleukemic K562 cells were transiently co-transfected with the pGL4.23 reporter construct and pGL-TK Renilla luciferase construct using Lonza 4D-Nucleofector system (Lonza). Firefly luciferase activity was measured 24 hours post-transfection by the Dual Luciferase Assay kit (Promega). Results were normalized to Renilla luciferase activity. These experiments were performed in triplicated for three independent times.

## Results

Two cohorts of children with newly-diagnosed ALL receiving 6-MP therapy were included in this study: the COG AALL03N1 and St. Jude Total XV clinical trials as the discovery and validation set, respectively (**Table 1**). After removing patients with known risk variants in *TPMT* and/or *NUDT15* and those without TGN measurements, 588 and 372 patients in the respective cohorts were included in the subsequent genetic association analyses.

**Table 1.**
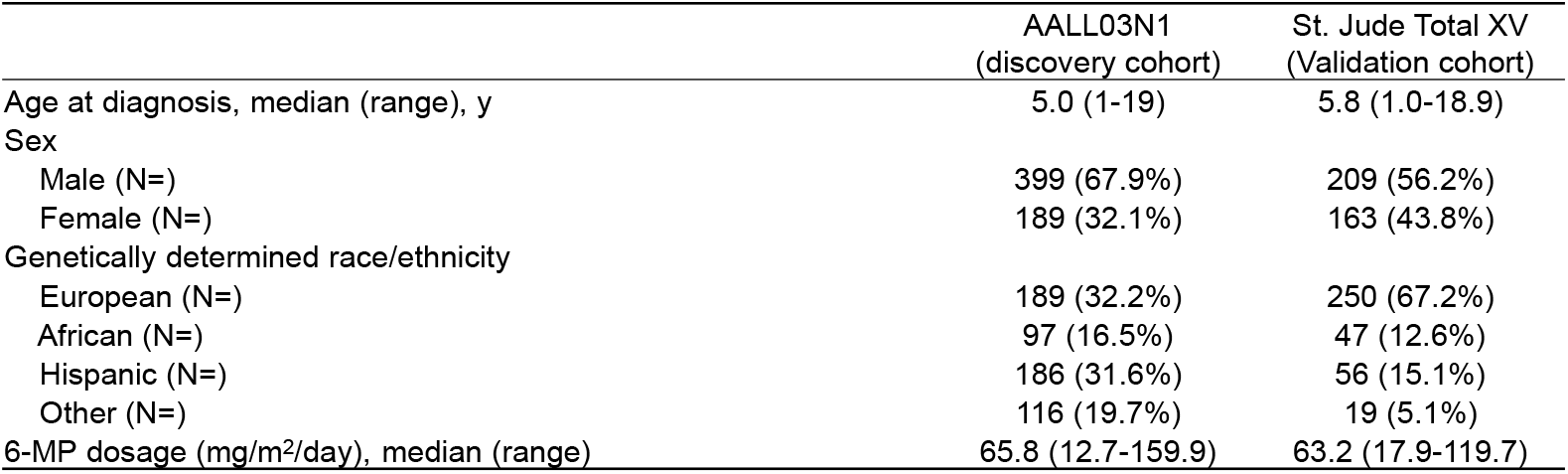
Patient characteristics of the COG AALL03N1 and the St. Jude Total XV cohorts, used for discovery and validation analyses, respectively.

|  | AALL03N1<br>(discovery cohort) | St. Jude Total XV<br>(Validation cohort) |
| --- | --- | --- |
| Age at diagnosis, median (range), y | 5.0 (1-19) | 5.8 (1.0-18.9) |
| Sex |  |  |
| Male (N=) | 399 (67.9%) | 209 (56.2%) |
| Female (N=) | 189 (32.1%) | 163 (43.8%) |
| Genetically determined race/ethnicity |  |  |
| European (N=) | 189 (32.2%) | 250 (67.2%) |
| African (N=) | 97 (16.5%) | 47 (12.6%) |
| Hispanic (N=) | 186 (31.6%) | 56 (15.1%) |
| Other (N=) | 116 (19.7%) | 19 (5.1%) |
| 6-MP dosage (mg/m<sup>2</sup>/day), median (range) | 65.8 (12.7-159.9) | 63.2 (17.9-119.7) |

### Targeted resequencing of *NT5C2* identified three missense variants

Targeted sequencing of *NT5C2* in the COG AALL03N1 cohort identified three missense variants, p.T3A (rs10883841), p.M15T (rs751896016), and I528V (rs756913240) (**Figure 1**). The p.T3A variant was recurrent with minor allele frequency of 9.3%, and p.M15T and I528V were observed only once in this cohort (**Figure 2A**). None of the three variants were predicted to be damaging by *in silico* predication tools SIFT or PolyPhen-2. The common variant p.T3A was not associated with TGN in patients (*P* > 0.05, **Figure 2A**), and the singleton variants were not evaluated in the association analysis due to low frequency. Furthermore, we characterized enzymatic properties of all three variant NT5C2 proteins with thiopurine metabolites as substrates. NT5C2 p.I528V showed a modest increase in 5’-nucleotidase activity against TIMP but not with TGMP (**Figure 2B**), whereas the other two variants showed a similar activity compared to wildtype NT5C2 (**Figure 2B and 2C**). Together, these results indicated that these three germline *NT5C2* missense variants did not affect erythrocyte TGN in children with ALL.

**Figure 1.**
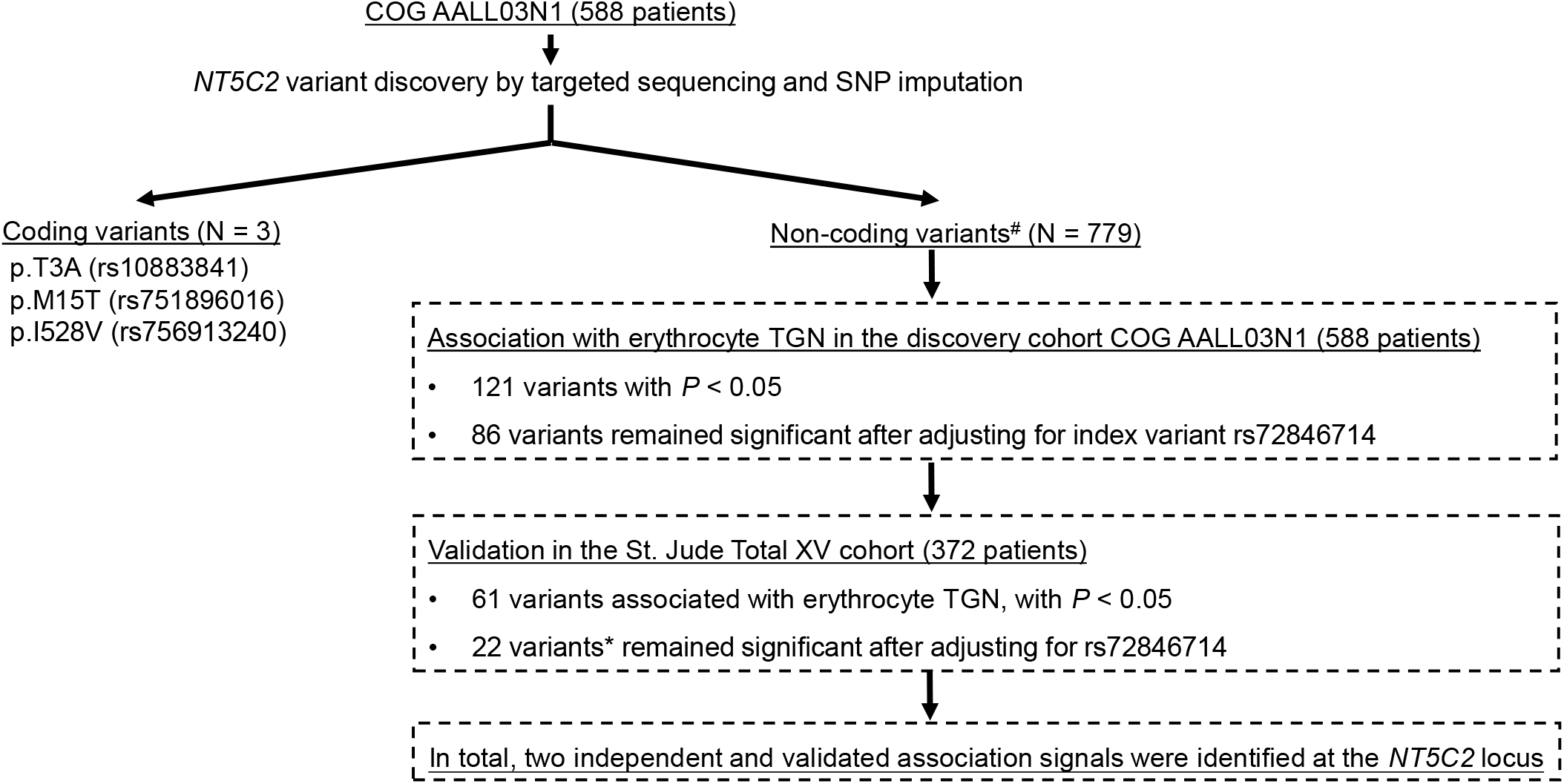
Discovery of *NT5C2* variants associated with 6-MP metabolism in children with ALL. Targeted sequencing coupled with SNP imputation in the discovery COG AALL03N1 cohort (588 patients) identified 782 variants at the *NT5C2* locus: three missense variants and 779 non-coding variants. The association of *NT5C2* SNP genotype with erythrocyte TGN was tested using a linear mixed-effects model with dose intensity and genetic ancestry as covariates. This analysis was restricted to patients with wildtype genotype for *TPMT* and *NUDT15*. To identify novel TGN associated variants, we also performed multivariate analysis conditioning on index variant rs72846714 identified from previous GWAS [34]. Validation was performed in the independent St. Jude Total XV cohort (372 patients), following the same analysis strategy. #non-coding variants with minor allele frequency<1% were excluded from the association analyses due to statistical consideration. * All 22 variants are in linkage disequilibrium with pair-wise LD r^2^ > 0.99.

**Figure 2.**
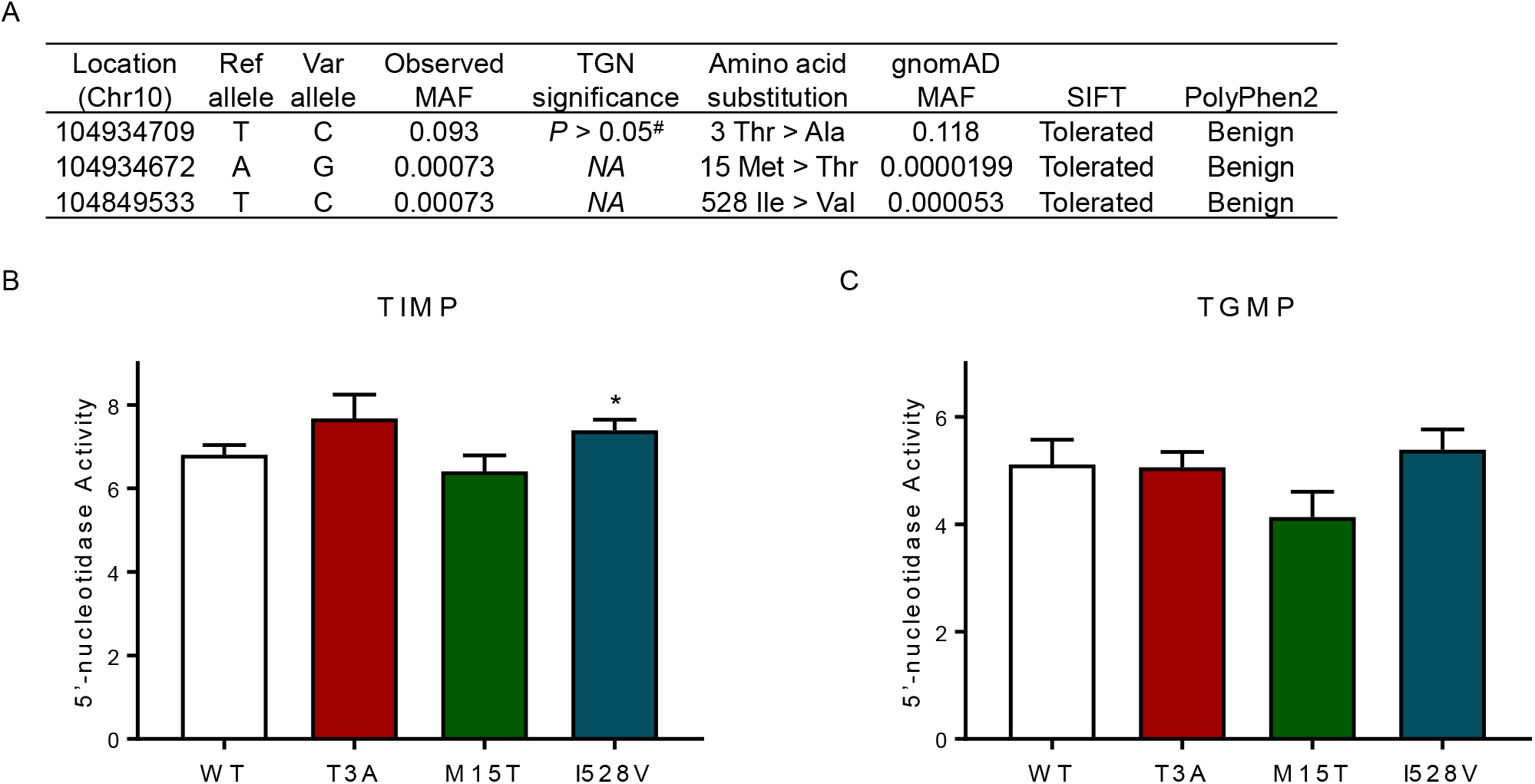
Functional characterization of *NT5C2* missense variants. (A) Targeted sequencing of *NT5C2* in the COG AALL03N1 cohort identified three missense variants. Chromosome location was aligned to GRCh37/hg19. SIFT and PolyPhen2 prediction tools were used to predict mutation function. MAF, minor allele frequency. (B) and (C) 5’-nucleotidase activity was determined of wildtype and variant NT5C2, with thiopurine metabolites as substrates. Purified NT5C2 proteins were incubated with TIMP or TGMP, and 5’-nucleotidase activity was measured by quantifying the release of free phosphate using the Malechite Green assay. Bars represent the mean values; the error bars represent the SD from triplicate. Ref allele, reference allele; Var allele, variant allele; #, *P* value of association of genotype and TGN was analyzed using a linear regression model with 6-MP dosage and genetic ancestry as covariates; *, *P* < 0.05 (Student *t* test). TGMP thioguanosine monophosphate; TIMP, thioinosine monophosphate.

### Association of non-coding variants in *NT5C2* with erythrocyte TGN

Non-coding variants in *NT5C2* were determined by targeted sequencing as well as SNP imputation. A total of 779 non-coding variants in the genomic region encompassing the *NT5C2* gene and the 200Kb flanking region with a minor allele frequency of at least 1%, were identified in the COG AALL03N1 cohort and tested for association with TGN (**Figure 1**). After adjusting for 6-MP dose and genetic ancestry as covariates,121 variants were nominally associated with erythrocyte TGN during 6-MP treatment (*P* < 0.05, **Figure 1, Figure S2 and Table S2**). These variants were then tested for replication in the St. Jude Total XV cohort, and 61 of 121 variants were validated for association with TGN at the *P* < 0.05 level (**Figure 1, Figure S2 and Table S2**), including rs72846714 which was reported in the previous GWAS of 6-MP metabolites [34] (**Figure 3A and 3B, Table S2**).

**Figure 3.**
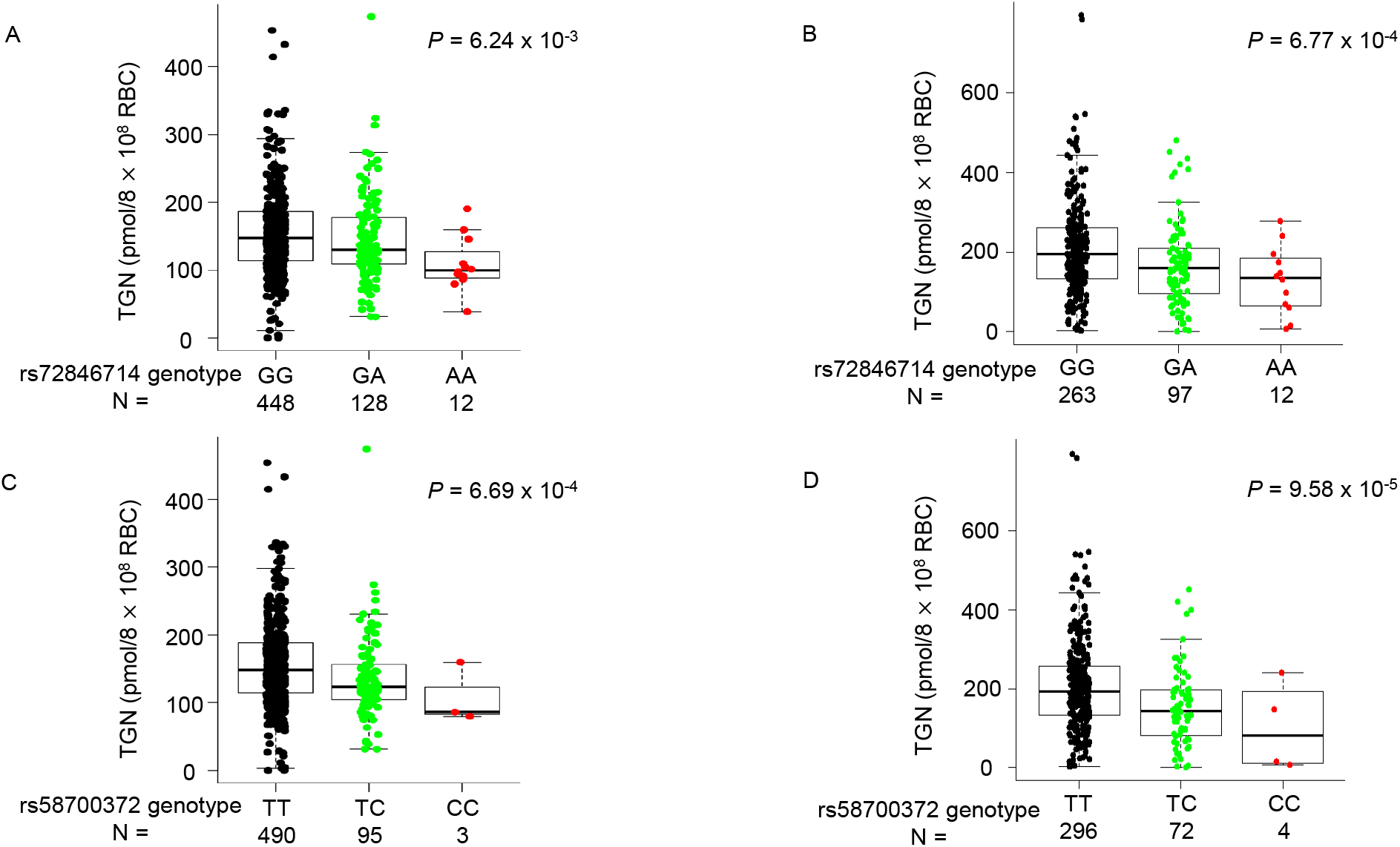
Association of non-coding *NT5C2* variants with erythrocyte TGN in the COG AAL03N1 and St. Jude Total XV cohorts. Effects of *NT5C2* SNP genotype on TGN were shown for two non-coding variants, rs72846714 and rs58700372, selected to represent two independent association signals at this locus. (A) and (B) TGN was significantly lower for patients heterozygous or homozygous for variant allele at rs72846714 in the COG AALL03N1 cohort (A) and the St. Jude Total XV cohort (B), compared with reference allele. Genotype-TGN association was presented in a similar format in (C) and (D) for rs58700372 in the COG AALL03N1 cohort (C) and the St. Jude Total XV cohort (D). *P* value was estimated using linear regression model with dosage and genetic ancestry as covariates. Each box included data between 25th and 75th percentiles, with horizontal line indicating median. Whiskers indicated maximal and minimal observations within 1.5x length of box.

Of 61 validated *NT5C2* SNPs, only 22 remained significant in both discovery and validation cohorts after adjusting for the GWAS hit rs72846714 [34]. This cluster of 22 variants were in strong linkage disequilibrium with each other (r^2^>0.99), and represented a novel association at the *NT5C2* locus independent of the GWAS signal. For example, in the discovery cohort COG AALL03N1, there was a significant decrease in TGN metabolite levels by genotypes at the independent SNP rs58700372 from wildtype (TT, 154.8 ± 61.1) to homozygotes (CC, 108.4 ± 44.4), with the heterozygotes (TC, 136.2 ± 61.1) exhibiting an intermediate level of TGN (*P* = 6.69 × 10^-4^; **Figure 3C**). This association was replicated in the St. Jude Total XV cohort: mean levels of the metabolite were 225.5 ± 132.9, 193.3 ± 86.0, and 165.5 ± 49.3 in wildtype, heterozygous and homozygous individuals, respectively (*P* = 9.58× 10^-5^. **Figure 3D**).

Interestingly, even within patients heterozygous for known *TPMT* and/or *NUDT15* variants, *NT5C2* genotype at rs72846714 and rs58700372 appeared to further distinguish patients with differential 6-MP metabolism although this only reached statistical significance in the St. Jude Total XV cohort (**Figure S3**). The frequency of variant allele at these two *NT5C2* SNPs also differ by race and ethnicity (**Figure S4**). Taken together, we identified two clusters of non-coding variants that independently influenced 6-MP metabolism, represented by rs72846714 and rs58700372.

### Functional exploration of the germline variants

To gain insight into the biological basis of the association signals at these non-coding *NT5C2* variants, we firstly examined chromatin accessibility inferred from the ATAC-seq of human hematopoietic cells and histone mark ChIP-seq signals form ENCODE project [42–44].

For the previous reported GWAS hit rs72846714, we found that it was in the intronic region with minimal overlap with regulatory elements (**Figure 4A**), indicating the lack of transcriptional regulation function of this variant. However, amongst variants in strong LD with rs72846714, we identified five SNPs that showed significant association with TGN in both discovery and validation cohorts and were also in proximity of open chromatin segments in multiple hematopoiesis cells (rs12261294, rs12240508, rs7091462, rs113642369, and rs12256506) (**Figure 4A**). We postulated that one of these variants may directly regulate *NT5C2* transcription and thus causal to the GWAS signal at this locus. We performed luciferase assay to test the effects of rs72846714 and each of the five LD variants on transcription activity and found that only rs12256506 substantially influenced enhancer activity (**Figure 4B**). The substitution of the reference allele with the variant allele at this variant resulted in a 1.48-fold increase in transcription activity (*P* = 0.0049, **Figure 4B**), consistent with higher expression of *NT5C2* and thus lower TGN in erythrocytes (**Figure 3A and 3B**). This is in line with the notion that NT5C2 degrades 6-MP active metabolites TIMP and TGMP, and thus its activity is inversely correlated with TGN.

**Figure 4.**
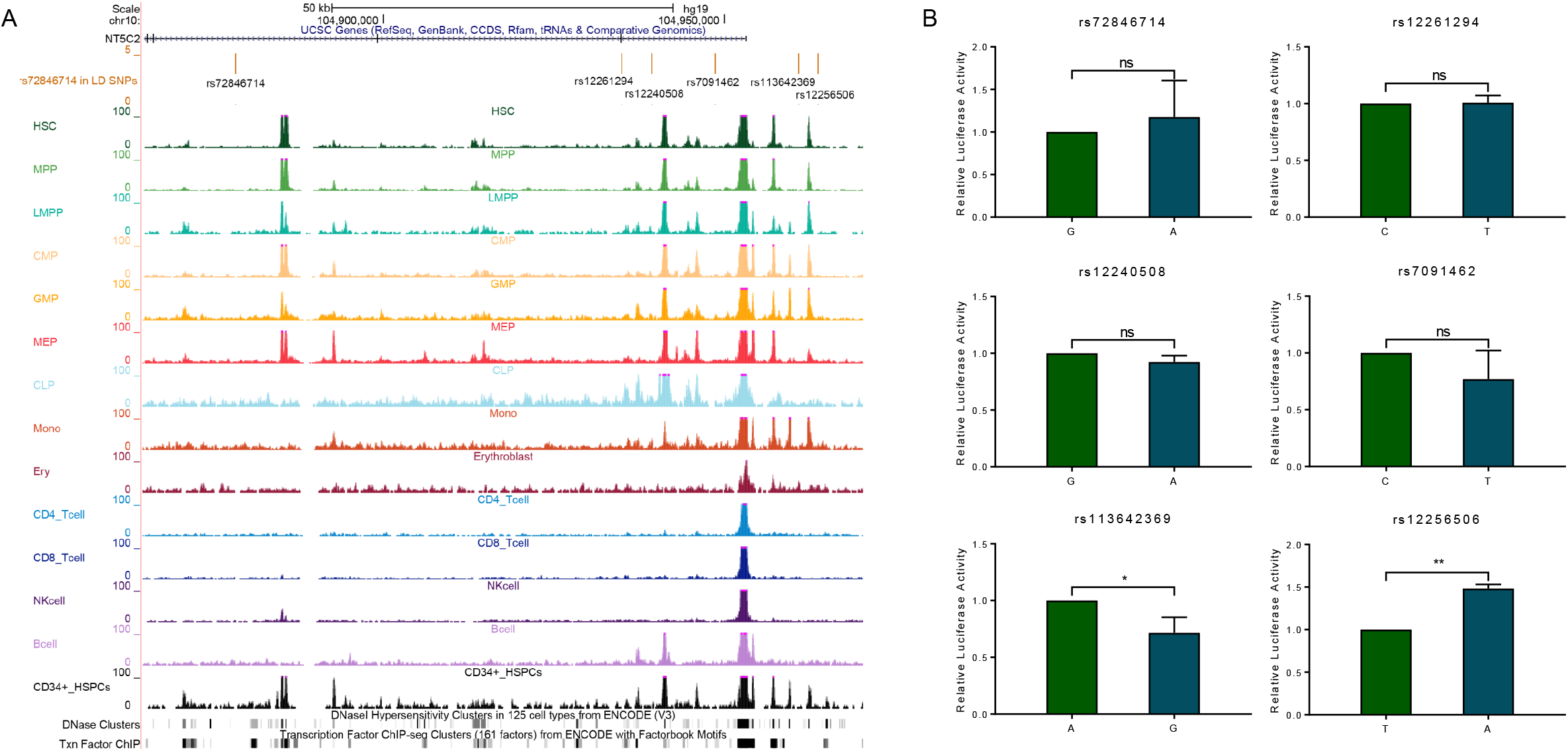
Functional characterization of *NT5C2* non-coding variant rs72846714 and its LD SNPs. A total of six SNPs were presented including rs72846714 and five SNPs that are in strong LD with the index variant (ALL cohorts, r^2^>0.6)) and also annotated to putative regulatory DNA. (A) Functional annotation of regulatory DNA at the *NT5C2* locus. Genomic positions, scale and gene structure for GRCh37/hg19 were shown on the top. rs72846714 and its LD SNPs were marked in the middle panel. ATAC-seq signals of hematopoietic cells, DNase and TF ChIP clusters were also included, using publicly available datasets [42–44]. (B) Enhancer activity of cis-elements encompassing these SNPs was tested using luciferase reporter assay in K562 cells. Relative luciferase activity indicated the ratio between the value of variant allele over that from the reference allele. All experiments were performed in triplicate and repeated three times. Statistical significance was evaluated by using two-sided Student’s *t* test. Error bars indicate SD. ns, not significant; *, *P* < 0.05; **, *P* < 0.01.

By contrast, the novel variant rs58700372 was located within an open chromatin region based on ATAC-seq signal in erythroblast cells (Figure 5A). DNase accessibility and transcription factors ChIP-seq also suggested active transcription in this region (Figure 5A). Using luciferase assay to test its effect on transcription activity, we found that substitution of the reference allele with the variant allele resulted in a 3.78-fold increase in transcription activity (Figure 5B). These findings indicate that rs58700372 directly potentiates the activity of an intronic enhancer, activates *NT5C2* transcription, and thereby reduces 6-MP metabolism.

**Figure 5.**
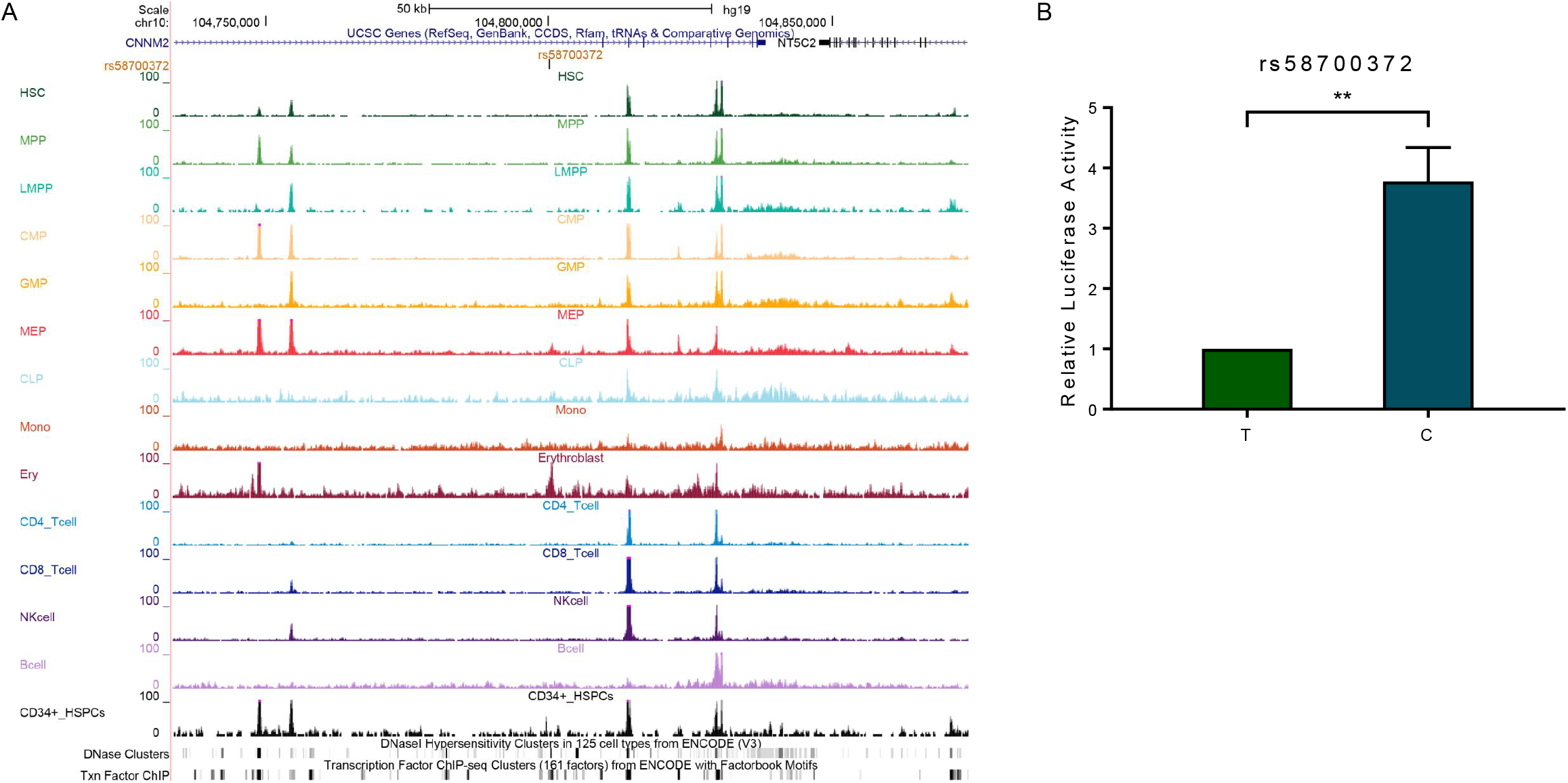
Functional characterization of *NT5C2* non-coding variant rs58700372. (A) Functional annotation of regulatory DNA within the genomic region encompassing rs58700372. Genomic positions, scale and gene structure for GRCh37/hg19 were shown on the top. rs58700372 was marked in the middle panel. ATAC-seq signals of hematopoietic cells, DNase and TF ChIP clusters were also included, using publicly available datasets [42–44]. (B) Luciferase assay was performed to determine enhancer activity of the DNA segment encompassing rs58700372 in K562 cells. Relative luciferase activity indicated the ratio between the value of the variant allele and that from the reference allele. This experiment was performed in triplicate and repeated three times. Statistical significance was evaluated by using two-sided Student’s *t* test. Error bars indicate SD; **, *P* < 0.01.

## Discussion

Prior pharmacogenetic studies have identified germline variants in *NUDT15* and *TPMT* associated with 6-MP sensitivity, and treatment modification based on *TPMT* and *NUDT15* genotype is a clear example of genetically guided precision medicine [22–27]. However, the majority of inter-individual variability in 6-MP pharmacokinetics remains unexplained. Recently, a genome-wide association study described a germline variant located in *NT5C2* intronic region associated with erythrocyte TGN during 6-MP therapy, pointing to potential effects of *NT5C2* germline variants on this phenotype [34].

NT5C2 is an ubiquitously expressed cytosolic nucleotidase involved in the maintenance of intracellular nucleotide pool homeostasis by promoting the clearance of excess purine nucleotides from cells [48, 49]. NT5C2 preferentially dephosphorylates the 6-hydroxypurine monophosphates inosine monophosphate (IMP), guanosine monophosphate (GMP), and xanthosine monophosphate (XMP), as well as the deoxyribose forms of IMP and GMP (dIMP and dGMP), facilitating the export of purine nucleosides. In the context of ALL therapy, NT5C2 can dephosphorylate and inactivate 6-MP metabolites TIMP and TGMP, and thus reduce the amount of TGN available for DNA incorporation and compromise cytotoxicity of this drug [28]. Acquisition of *NT5C2* mutations in relapsed ALL impedes the formation of DNA-TGN and is a frequent mechanism of 6-MP drug resistance [29–32].

In this study, we comprehensively evaluated *NT5C2* germline variants for their association with the 6-MP metabolite TGN during ALL therapy. We identified 61 *NT5C2* variants significantly associated with TGN after adjusting for dose and genetic ancestry in both discovery and validation cohorts. In line with previous GWAS [34], the variant A allele of rs72846714 was associated with lower TGN levels, indicating that this pharmacogenetic association is robust across different chemotherapy regimens and different patient populations. Importantly, we also identified a novel variant, namely rs58700372, that exhibited independent effect on 6-MP metabolite beyond that of the GWAS hit, suggesting multiple functional variants at this locus. However, neither of these two variants were associated with 6-MP dose intensity in our analyses (**Figure S5**), plausibly because these germline *NT5C2* SNPs only modestly alter drug metabolism to a degree that is insufficient to cause clinical toxicity or drug resistance. It is also worth noting that there was a trend for *NT5C2* variants to be associated with TGN even within patients hemizygously deficient of *TPMT* and/or *NUDT15* (**Figure S3**). This is relevant because these individuals exhibit wide variability in 6-MP tolerance and additional biomarkers are needed to precisely predict thiopurine toxicity. It remains possible that many other genetic variants influence 6-MP metabolism with relatively low penetrance, and their combined effects may warrant clinical consideration. There was also a trend that the variant alleles at rs72846714 and rs58700372 were linked to lower MeTIMP in erythrocytes in our ALL cohorts but it did not reach statistical significance (**Figure S6**).

Unlike *TPMT* or *NUDT15*, there are no common germline coding variants in *NT5C2* that affect its activity. The majority of pharmacogenetic variants identified in this gene are non-coding. Located in intronic and intergenic regions, these *NT5C2* variants primarily affect gene function by transcription regulation. For example, the association signal at the previously reported GWAS hit rs72846714 was likely explained by a proximal variant rs12256506 that activates *NT5C2* transcription in-cis. Our analysis also identified rs58700372 for its independent association with TGN beyond that of the GWAS SNP. Our functional study indicated that this variant directly alters the activity of an intronic enhancer, with the variant allele linked to higher transcription activity and reduced 6-MP activation (lower TGN). Of note, rs58700372 is located ~50kb downstream of the coding region of *NT5C2*, and the potential mechanism by which this variant regulates *NT5C2* transcription may involve chromatin remodeling and long-distance interaction, which need to be examined in future studies. It remains unclear whether functional effects of these two *NT5C2* SNPs are restricted to erythrocytes and future studies are warranted to explore cellular models of a broader range of tissue origins.

In conclusion, we comprehensively analyzed the role of *NT5C2* germline variants in 6-MP metabolism during ALL therapy and identified functional SNPs that independently and directly regulate NT5C2 activity and 6-MP metabolism. Our results indicate that *NT5C2* germline variation contributes to inter-patient variability in thiopurine drug disposition.

### Study Highlights

#### What is the current knowledge on the topic?

Levels of intracellular 6-MP metabolites can inform drug toxicity and response and potentially inform treatment individualization. Recent GWAS study reported initial evidence linking *NT5C2* germline variants to 6-MP metabolism. However, the full spectrum of genetic polymorphism in *NT5C2 and* their impact on inter-patient variability in thiopurine drug disposition are largely unknown.

#### What question did this study address?

This pharmacogenomics study comprehensively evaluated the contribution of genetic variations in *NT5C2* to 6-MP metabolite TGN in two ALL cohorts and identified two representative SNPs that independently affected 6-MP metabolism. Importantly, we also sought to understand the molecular mechanism by which these non-coding variants modulate NT5C2 function.

#### What does this study add to our knowledge?

This is one of the first studies to describe the full spectrum of genetic variants in *NT5C2* and examined their effects on thiopurine metabolism in large cohorts of patients. We also provided functional evidence of how coding and non-coding variants modulate NT5C2 activity. Our results firmly established the importance of *NT5C2* germline variation in explaining inter-patient variability in thiopurine drug disposition.

#### How might this change clinical pharmacology or translational science?

This study identified novel genetic variants significantly associated with thiopurine drug activation, therefore expanding the list of pharmacogenetic factors that may aid the optimization of pharmacotherapy using this class of drugs.

## Acknowledgment

We thank the patients and parents who participated in the clinical studies included in this report. This work was partially supported by National Institutes of Health grants GM118578, GM115279, and CA096670.

## Author contributions

C.J., H.Z. and J.J.Y. wrote the manuscript; J.J.Y. designed the research; C.J., Z.P.L. and T.M. performed the research; W.J.Y., C.S., W.T.Y., M.X.Q and M.T. analyzed the data; C.C.L., C.H.P., W.E., S.B., K. S., and K.R.C. and J.J.Y. contributed new reagents and/or analytical tools.

## Supplementary Information Titles

Figure S1.

Figure S2.

Figure S3.

Figure S4.

Figure S5.

Figure S6.

Table S1. Primers used in this study.

Table S2. TGN significantly associated *NT5C2* SNPs in COG AALL03N1 cohort.

