## Supplementary Figures for "Effects of *NT5C2* germline variants on 6-mecaptopurine metabolism in children with acute lymphoblastic leukemia"

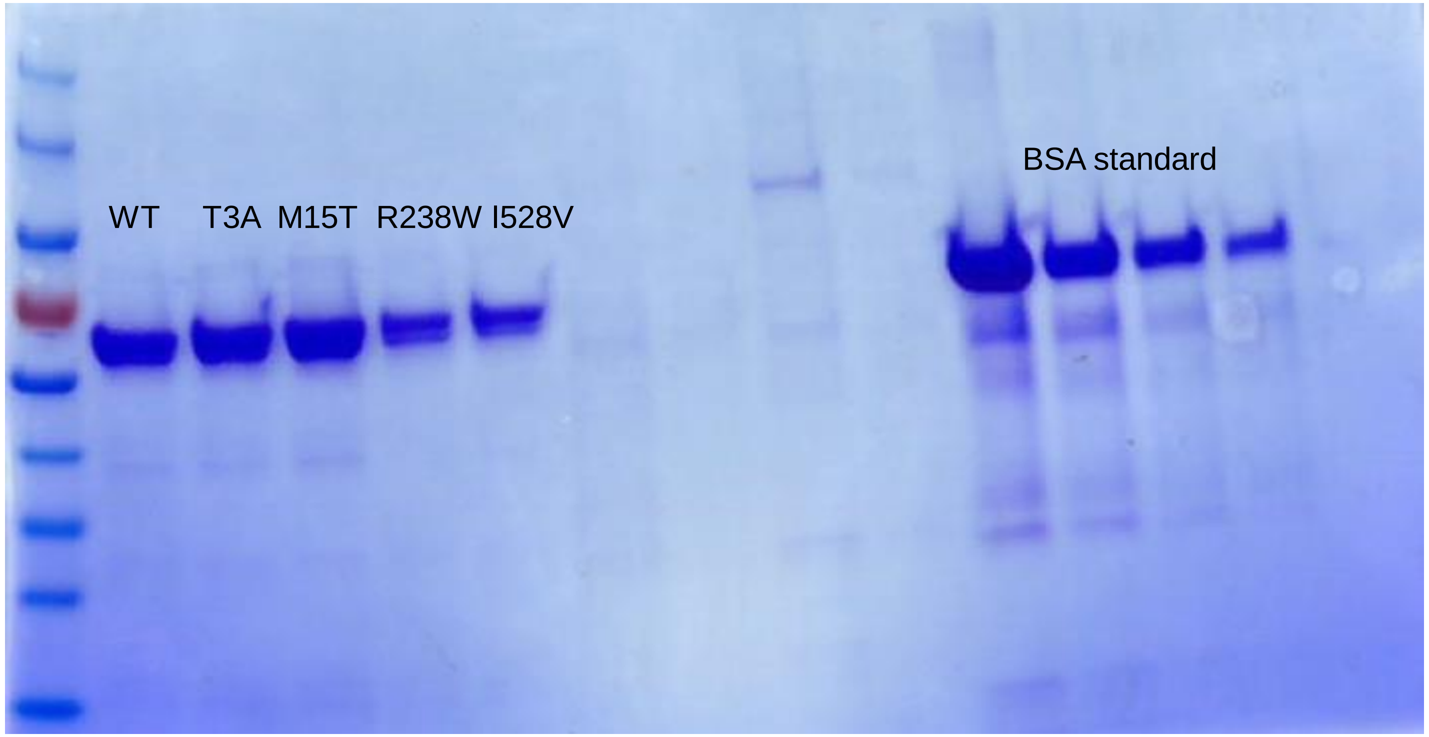


**Figure S1. Coomassie brilliant blue staining of NT5C2 proteins.** Right left lane was protein maker; Left second to sixth lanes were NT5C2 wildtype and mutant proteins(NT5C2 wt, T3A, M15T, R238W and I528V); Four lanes right side were BSA standard.

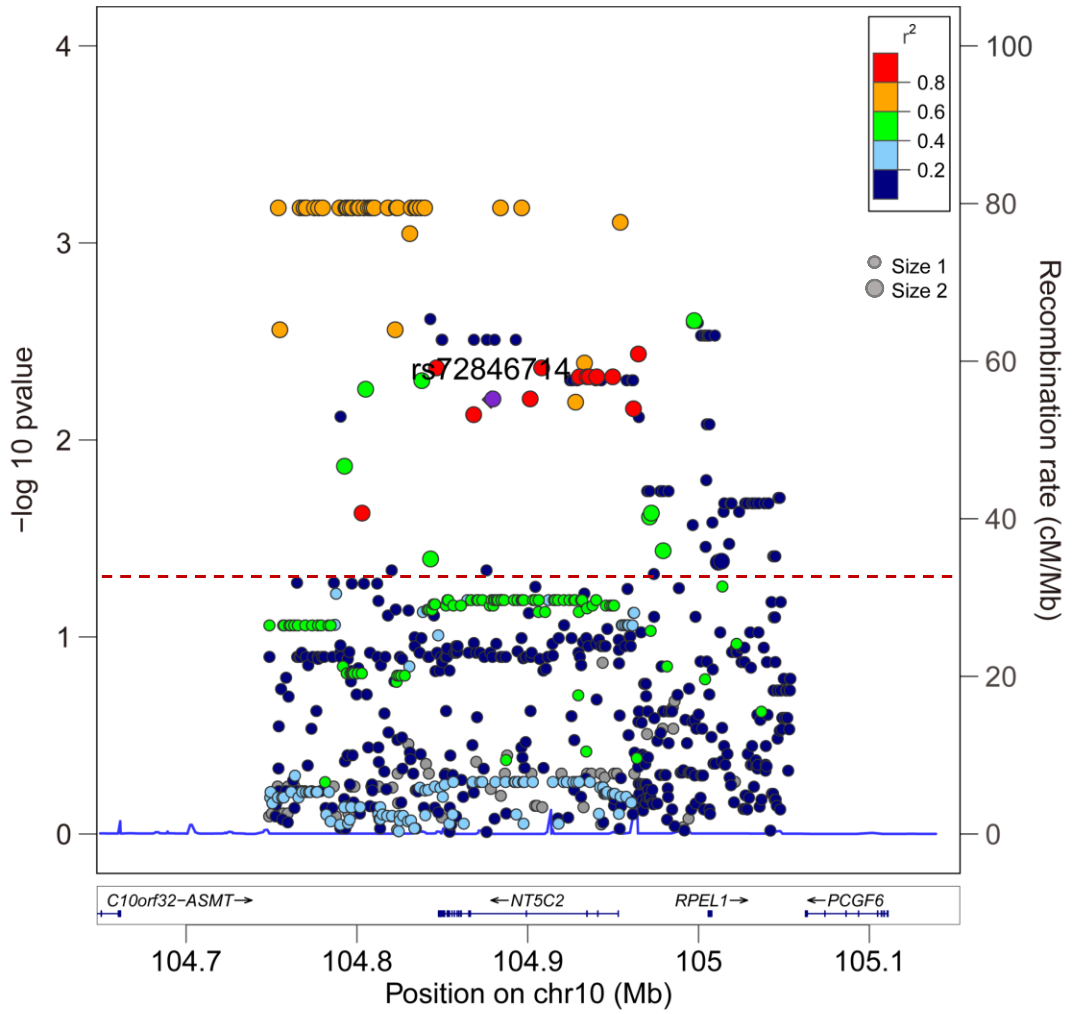


**Figure S2. Locus Zoom plot for 100kb upstream and downstream of *NT5C2* gene loci.** The negative logarithm of *P* values (y axis) were plotted for each single nucleotide polymorphism (SNP) in COG AALL03N1 cohort. The red horizontal line indicated the significant threshold (*P* = 0.05). Circus dot with Size 1 denoted 779 noncoding variants which were identified in COG AALL03N1 cohort, and circus dot with Size 2 represented significant variants in both COG AALL03N1 and St. Jude Total XV cohort. Purple dot (chr10:104878454) was rs72846714. LD score (as indicated by colors) was calculated LD with rs72846714 as index SNP. Genomic positions and gene structure for the human genome assembly February 2009 (GRCh37/hg19) were shown on the bottom.

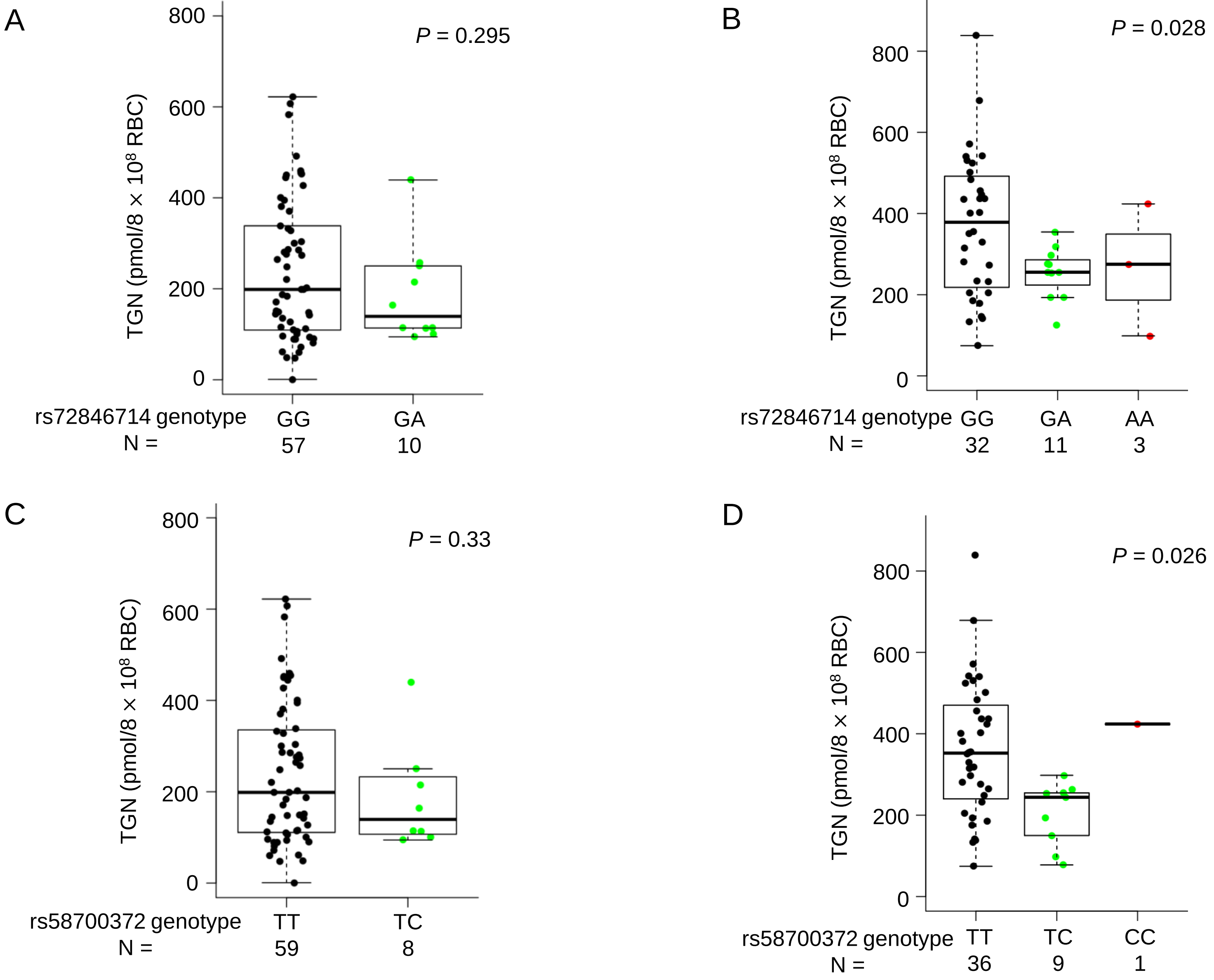


**Figure S3. Association of rs72846714 and rs58700372 with erythrocyte TGN in *TPMT* and/or *NUDT15* heterozygous patients in the COG AAL03N1 and St. Jude Total XV cohorts.** Effects of *NT5C2* SNP genotype on TGN were shown for two non-coding variants, rs72846714 and rs58700372 in patients with *TPMT* and/or *NUDT15* heterozygous genotype. . (A) and (B) TGN was lower for patients heterozygous or homozygous for variant allele at rs72846714 in the COG AALL03N1 cohort (A) and the St. Jude Total XV cohort (B), compared with reference allele, but the differences were only significant in St. Jude Total XV cohort. Genotype-TGN association was presented in a similar format in (C) and (D) for rs58700372 in the COG AALL03N1 cohort (C) and the St. Jude Total XV cohort (D). *P* value was estimated using linear regression model as described. Each box included data between 25th and 75th percentiles, with horizontal line indicating median. Whiskers indicated maximal and minimal observations within 1.5x length of box.

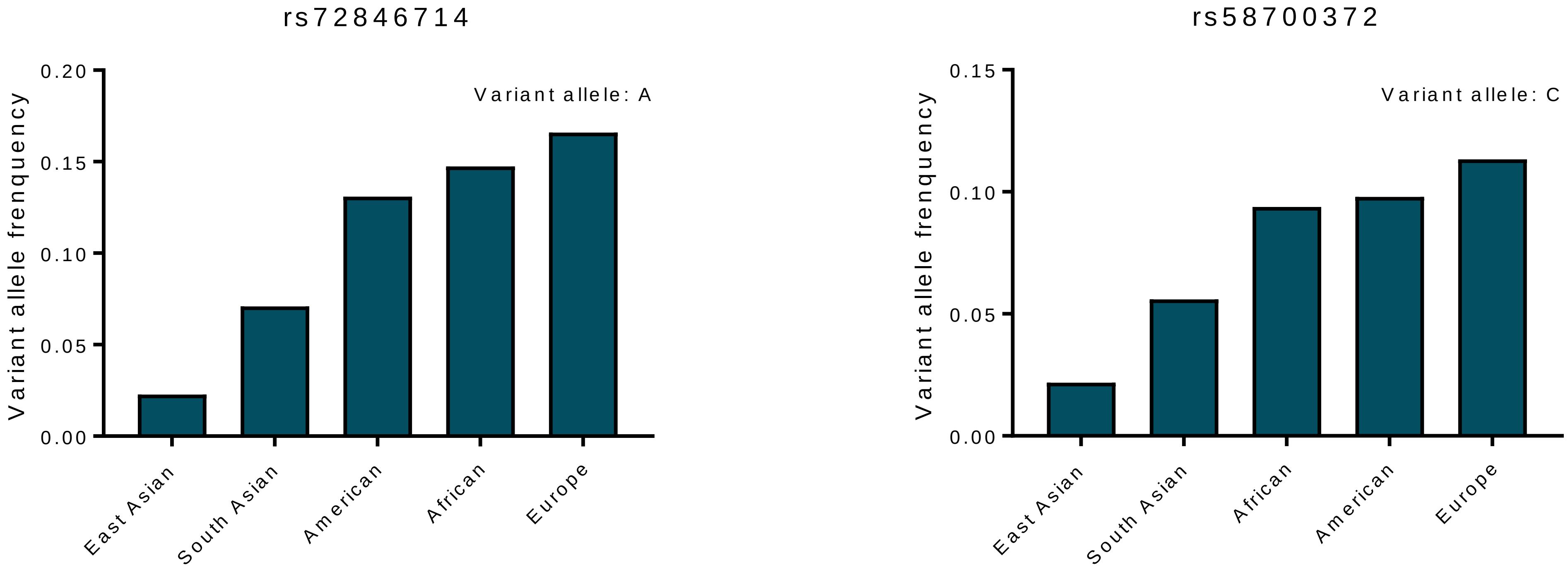


**Figure S4. Variant allele frequency of rs72846714 and rs58700372 in different ethnic groups.** Allele frequency data was retrieved from 1000 Genomes Project.

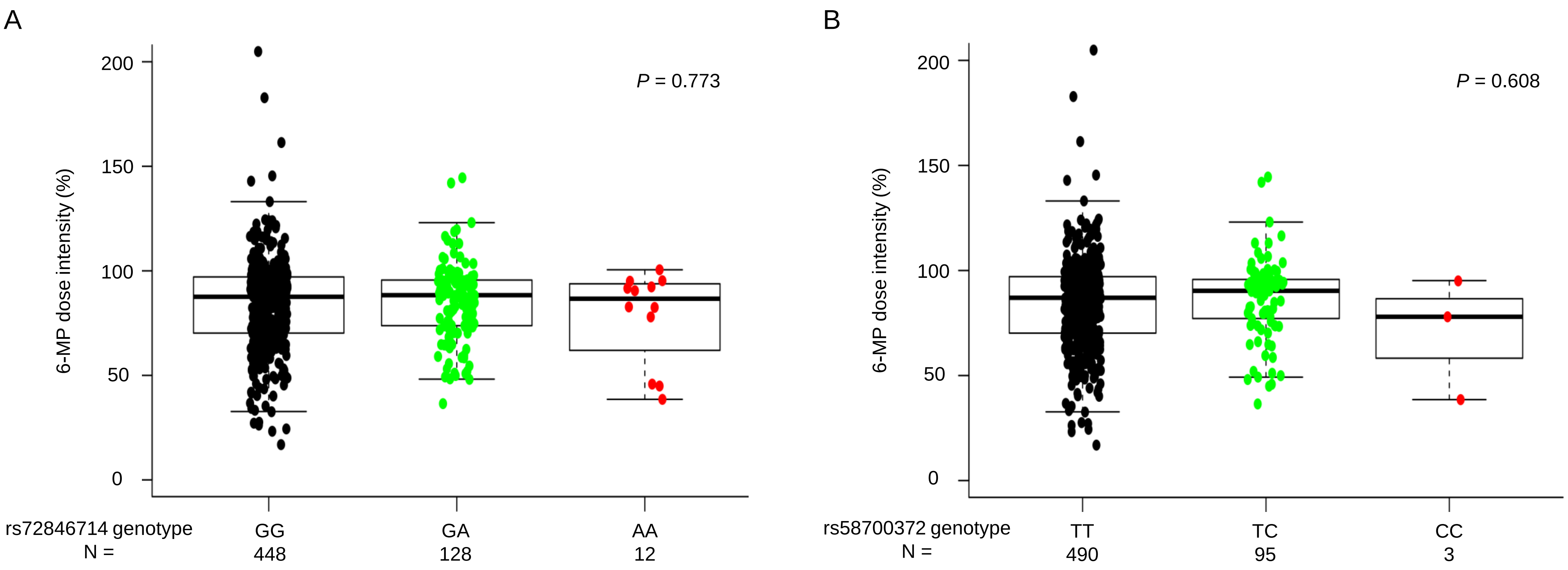


**Figure S5. Association of non-coding *NT5C2* variants with 6-MP dose intensity in the COG AAL03N1 cohort.** For patients enrolled onto COG AALL03N1 protocol, 6-MP dose was adjusted during maintenance therapy on the basis of host toxicities (myelosuppression and infections), and dose intensity was defined as ratio of prescribed dose over protocol planned dose (75 mg/mg$^2$ per day). Dose intensity was measured longitudinally over 6 months and was shown as single cumulative value for study period. Effects of *NT5C2* SNP genotype on 6-MP dose intensity were shown for two non-coding variants, rs72846714 (A) and rs58700372 (B). *P* value was estimated using linear regression model as described. Each box included data between 25th and 75th percentiles, with horizontal line indicating median. Whiskers indicated maximal and minimal observations within 1.5x length of box.

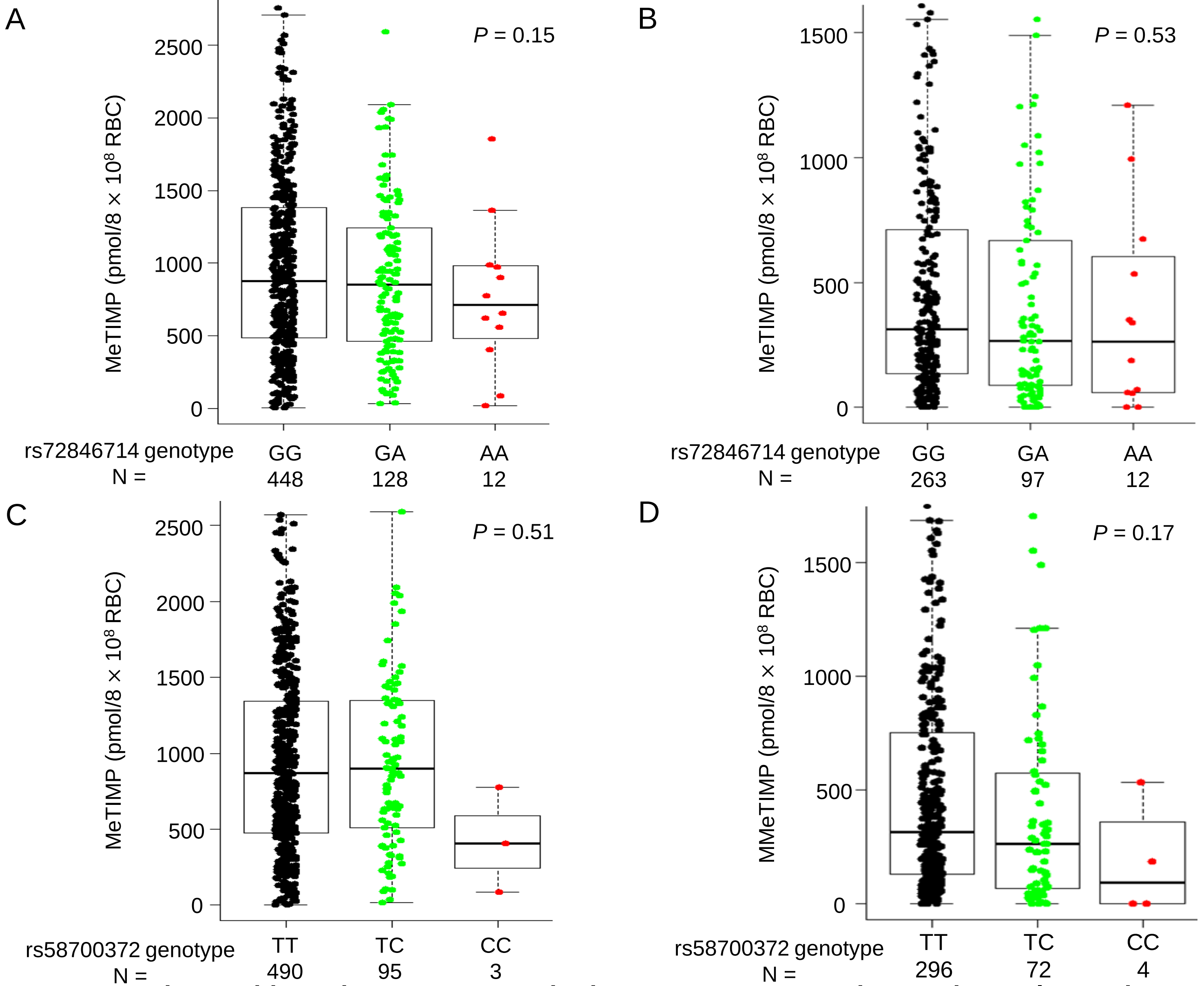


**Figure S6. Association of non-coding *NT5C2* variants with erythrocyte MeTIMP in the COG AAL03N1 and St. Jude Total XV cohorts.** Effects of *NT5C2* SNP genotype on MeTIMP were shown for two non-coding variants, rs72846714 and rs58700372, selected to represent two independent TGN-association signals at this locus. (A) and (B) MeTIMP was lower for patients heterozygous or homozygous for variant allele at rs72846714 in the COG AALL03N1 cohort (A) and the St. Jude Total XV cohort (B), compared with reference allele, but the differences were not significant. Genotype-MeTIMP association was presented in a similar format in (C) and (D) for rs58700372 in the COG AALL03N1 cohort (C) and the St. Jude Total XV cohort (D). *P* value was estimated using linear regression model as described. Each box included data between 25th and 75th percentiles, with horizontal line indicating median. Whiskers indicated maximal and minimal observations within 1.5x length of box.
